# Hepatic cellular stress response pathways exhibit species differences in basal and inducible activity

**DOI:** 10.1101/2025.09.22.677731

**Authors:** Hannah Coghlan, Dominic Williams, Andrew R. Jones, Sophie Regan, Bhavik Chouhan, Ian M. Copple

## Abstract

Cellular stress response pathways such as the NRF2 oxidative stress response, endoplasmic reticulum (ER) stress response and macroautophagy afford protection against many forms of drug toxicity, including the liver toxicity associated with the formation of reactive drug metabolites. To maximise the translatability of preclinical toxicology studies, an understanding of the relative hepatic stress response capacities of humans and widely-used preclinical animal species is vital. In control liver tissue, the basal gene and protein expression of stress response pathway components was found to be greater in rodents than non-rodent preclinical species and humans. In addition, following in vitro exposure to pharmacological modulators of the NRF2 and ER stress responses, rodent hepatocytes generally displayed a greater capacity, relative to those of non-rodent preclinical species and humans, for adaptation to cellular stress. Consistent with the reported lower concordance of drug toxicity between humans and rats, the latter displayed a greater level of Torin1-induced autophagic flux than all other species, while the robust transcriptional responses to thapsigargin-induced endoplasmic reticulum stress and Bardoxolone- or Ki696-mediated NRF2 activation were comparable between mouse and rat hepatocytes. In all, our results indicate that rodent preclinical species possess a greater basal and adaptive hepatic capacity for mitigation of chemical insult than non-rodent preclinical species and humans. This study represents the first to provide a comprehensive comparison of stress response pathway capacity of humans and the animal species most commonly used for preclinical drug safety assessment. Our findings can be used to inform the selection of species for safety testing of drugs with a liability for reactive metabolite-mediated liver toxicity.

## Introduction

There is a clear aspiration that the drug discovery and development process will transition away from a reliance on *in vivo* animal studies, due to ethical and translational concerns (Wadman, 2023, U.S. Food and Drug Administration, 2025). Yet, at present, almost all drug development programmes use *in vivo* animal studies to assess the toxicity liabilities of a new drug (Sewell *et al*., 2024). Despite this, animal studies predict human drug-induced liver injury (DILI) only marginally more accurately than random chance (55-57% predictive rate) (Olson *et al*., 2000, Tamaki *et al*., 2013).

Differences in the activity, specificity and tissue distribution of drug metabolising enzymes (Baillie and Rettie, 2011) and transporter proteins (Wang *et al*., 2015), drug target structure and expression (Eastwood *et al*., 2010), and immunology (Bjornson-Hooper *et al*., 2022) between humans and preclinical species have previously been shown to affect species sensitivity to drug toxicity. For example, preclinical detection of the mitotoxic potential of fialuridine was impeded by species-specific differences in subcellular localisation of the equilibrative nucleoside transporter 1, which resulted in fatal human DILI in clinical trials (McKenzie *et al*., 1995, Lee *et al*., 2006). Additionally, the antidiabetic drug troglitazone was withdrawn from the market due to severe clinical hepatotoxicity that was not predicted during preclinical toxicology studies due to species-specific differences in drug exposure and clearance (Smith, 2003, Shen *et al*., 2012).

Many hepatotoxins exert their effects via chemically reactive metabolites (CRMs), which cause widespread cellular injury via non-specific covalent interactions with macromolecules including nucleic acids, proteins, and lipid bilayers (Attia, 2010). Several stress responses, including the nuclear factor erythroid 2-related factor (NRF2)-mediated response to oxidative stress and unfolded protein response (UPR) to endoplasmic reticulum (ER) stress, as well as autophagy, have been shown to afford protection against the DILI caused by many drugs and their CRMs, including acetaminophen (APAP), diclofenac, amiodarone and isoniazid (Fredriksson *et al*., 2014, Liu *et al*., 2013, Wandrer *et al*., 2020, Zhang *et al*., 2016). Therefore, differences between humans and preclinical animal species in the capacity to respond to and ameliorate chemical insult via these pathways could contribute to a species’ relative sensitivity to certain forms of DILI.

We recently reported that, upon challenge with equivalent chemical insult resulting from administration of APAP, the rat exhibits a greater basal and adaptive capacity for hepatic stress responses compared to the mouse (Russomanno *et al*., 2023). However, the true extent of inter-species differences in stress response capacity is relatively unknown, with current literature focussing only on rodent preclinical species and lacking a human comparator, which is necessary to establish translatability. To address this knowledge gap, the present study utilised liver tissue and primary hepatocytes from humans – as the benchmark species – and a broader range of relevant preclinical species to assess inter-species differences in hepatic stress response capacity. Our results show that rodent preclinical species possess a generally greater capacity for stress response induction than humans and non-rodent preclinical species, which may afford these species reduced sensitivity to CRM-mediated DILI. As such, these results highlight a major limitation of preclinical animal species and reinforce the need for reduced reliance on these models.

## Methods

Unless otherwise stated, materials were purchased from Thermo Fisher Scientific (Waltham, MA, USA). Torin1, bafilomycin A1 (BA1) and thapsigargin were purchased from Selleckchem (Planegg, Germany), while bardoxolone methyl ester (CDDO-Me) and Ki696 were purchased from Sigma (St. Louis, MO, USA). Bioinformatic analyses were conducted in RStudio using R version 4.4.1.

### Liver tissue

Human liver tissue samples from patients undergoing planned liver resections were obtained, with full written, informed consent, by qualified medical staff at Aintree University Hospital (Liverpool, UK) and stored at -80 °C until use. The study protocol was approved by the National Health Service North West Liverpool Central Research Ethics Committee (11/NW/0327) and adhered to the 1975

Declaration of Helsinki. Liver tissues from untreated rodents were kindly provided by members of research groups within the University of Liverpool (Liverpool, UK). Liver tissues from vehicle-treated cynomolgus macaques and beagle dogs were provided by Charles River Laboratories (Harlow, UK). Full details of the patients and animals from which tissue samples were obtained are presented in Supplementary Table 1.

### Primary hepatocyte culture

Cryopreserved primary human, cynomolgus macaque and beagle dog hepatocytes were obtained from Primacyt (Schwerin, Germany). CD1 mouse and Sprague-Dawley rat CryostaX hepatocytes were obtained from SEKISUI Xenotech (Kansas City, KS, USA). To minimise the effect of differential hepatocyte culture procedures on stress response activity, cells were thawed, plated and cultured according to the same protocol. Specifically, cells were cultured in Corning BioCoat Collagen I-coated plates (Corning, Flintshire, UK) in Williams Medium E supplemented with 2 mM L-glutamine (Sigma), 100 nM dexamethasone (Sigma), 1X insulin-transferrin-selenium and 100 U/mL penicillin/100 µg/mL streptomycin (Sigma). For all experiments, media was also supplemented with 10% (v/v) fetal bovine serum (FBS) to enable to proper autophagic function (González-Rodríguez *et al*., 2014, Spormann *et al*., 2020) and to prevent aberrant basal activation of autophagy affecting other stress response pathways with which it is integrated (Kalinin *et al*., 2023). Culture medium was replaced at 6 h and 24 h post-plating, at which time cells were exposed to positive control modulators of the stress response pathways under investigation, or 0.5% (v/v) DMSO, for 2-24 h. Hepatocyte morphology was monitored by light microscopy to ensure the retention of differentiation throughout the experiment (Supplementary Figure 1).

### Western blotting

Briefly, radioimmunoprecipitation assay buffer (Sigma) was used for snap-frozen liver tissue homogenisation and hepatocyte lysis. Western blotting was performed as described by Russomanno *et al*. (2023). Supplementary Table 2 details the antibodies used in this study. NCBI Protein BLAST (Camacho *et al*., 2009) was used to confirm that the immunogen of each antibody displayed high homology (>85%) with the target protein of each species of interest, indicating a comparable antibody affinity across species (Supplementary Table 3). Protein expression was quantified using Image Lab version 6.1 (Bio-Rad). Target protein normalisation was performed using β-actin or total protein expression (Ponceau S signal).

### RNA-seq

RNA was isolated from hepatocytes using the Monarch Total RNA Miniprep Kit (New England Biolabs, MA, USA) in accordance with the manufacturer’s protocol, including an on-column DNase digestion. RNA-seq analysis was performed by Eurofins Genomics LLC (Konstanz, Germany) and reads aligned to the appropriate reference genome. As low read count genes are less accurately quantified (Islam *et al*., 2014), genes with fewer than 30 reads per kilobase per million mapped reads (RPKM) in all samples were excluded. Differential gene expression analysis was performed using the DESeq2 package (version 1.44.0) (Love *et al*., 2014). Genes were considered to be significantly changed when a false discovery rate-adjusted *p*-value (*p*_adj_) ≤ 0.05 and fold change (FC) ≤ -1.5 or ≥ 1.5 were achieved. Heatmaps were generated using the pheatmap package (version 1.0.12) (Kolde, 2019).

Gene set enrichment analysis (GSEA) was performed using the Bioconductor package clusterProfiler (version 4.12.6) (Wu *et al*., 2021, Yu *et al*., 2012). Genome-wide annotation of human, mouse, rat and dog genes was performed using the packages ‘org.Hs.eg.db’ (version 3.19.1), ‘org.Mm.eg.db’ (version 3.19.1), ‘org.Rn.eg.db’ (version 3.19.1) and ‘org.Cf.eg.db’ (version 3.19.1), respectively (Carlson, 2019a, Carlson, 2019b, Carlson, 2019d, Carlson, 2024). As no package was available for annotation of cynomolgus macaque genes, the rhesus macaque (‘org.Mmu.eg.db’, version 3.19.1) package was used instead (Carlson, 2019c). Gene ontology (GO) biological processes with *p* ≤ 0.05 and normalised enrichment score (NES) ≤ -1.5 or ≥ 1.5 were considered to be significantly enriched or underrepresented, respectively. Unadjusted *p*-values were reported from GSEA to enable exploration of potentially relevant biological effects of treatments that elicited subtle transcriptional responses in some species, which are obscured by stringent multiple comparison corrections. The Bioconductor package rrvgo (version 1.16.0) was used to reduce GO term redundancy into encompassing parent terms based on semantic similarity (Sayols, 2020).

### Analysis of public RNA-seq data

Basal hepatic RNA-seq datasets for CD1 mice, Sprague-Dawley rats, cynomolgus macaques and beagle dogs (n=3/sex) were obtained from Krause *et al*. (2023) (GEO: GSE219045). Human hepatic RNA-seq data (n=262) was obtained from the GTEx Portal on 10^th^ June 2025 (Stanfill and Cao, 2021).

One-to-one human orthologs were mapped onto each species’ gene set using Ensembl BioMart (Dyer *et al*., 2024). The resulting datasets were combined to yield a final set of 11,074 protein-coding genes with one-to-one orthologs in all species of interest. This gene set was then normalised via ranking mean RPKM to enable inter-species comparison. This dataset was filtered by a lists of stress response-associated genes obtained from Qiagen Ingenuity Pathway Analysis (IPA) (Krämer *et al*., 2013).

### Statistical analysis and data presentation

Statistical analyses were performed in GraphPad Prism 10. Normal distribution of data was assessed using a Shapiro-Wilk test. Normally distributed data was analysed using a 2-tailed unpaired Student’s *t*-test or One-Way ANOVA followed by Tukey’s post-hoc test, as appropriate. Where data where non- normally distributed, a Mann-Whitey U test or Kruskal-Wallis test followed by Dunn’s post-hoc test was used, as appropriate. Inter-species comparisons were performed on ranked basal RNA-seq data using a Friedman test followed by a Conover-Iman post-hoc test. Unless otherwise stated, a statistical significance threshold (*p* or *p*_adj_) of 0.05 was utilised. Where gene or protein expression was compared between species, the human name (e.g., *NQO1* or NQO1) was used in figures and description of the results.

## Results

### SPECIES DIFFERENCES IN BASAL HEPATIC STRESS RESPONSE PATHWAY ACTIVITY

To assess differences in a species’ capacity to detoxify chemical insult and maintain homeostasis under basal conditions, public liver RNA-seq data - obtained from the human GTEx database (Stanfill and Cao, 2021) and the preclinical species gene expression database (Krause *et al*., 2023) – were leveraged to assess the basal hepatic expression of panels of stress response-associated genes (Figure 1A). No significant inter-species differences in the basal hepatic expression of NRF2-mediated oxidative stress response, UPR or autophagy genes were identified, although the mean rank of expression was generally greater in rodents than in non-rodents. To further investigate this, western blotting was used to measure the expression of key stress response components, selected from existing literature, in untreated liver tissues (Figure 1B). Humans generally displayed the lowest basal hepatic expression of these proteins of all the species investigated (significantly lower expression than at least one preclinical species for 8/9 proteins investigated) (Figure 1C). In contrast, rodents tended to display the greatest expression of stress response components amongst the preclinical species assessed (Figure 1C). Overall, the lower expression of key stress response components in human liver may indicate that this species posesses a weaker basal ability to mitigate hepatic chemical insult than the preclinical species investigated while, conversely, preclinical rodent species may possess a relatively enhanced basal capacity for detoxifying chemical insult.

**Figure 1.**
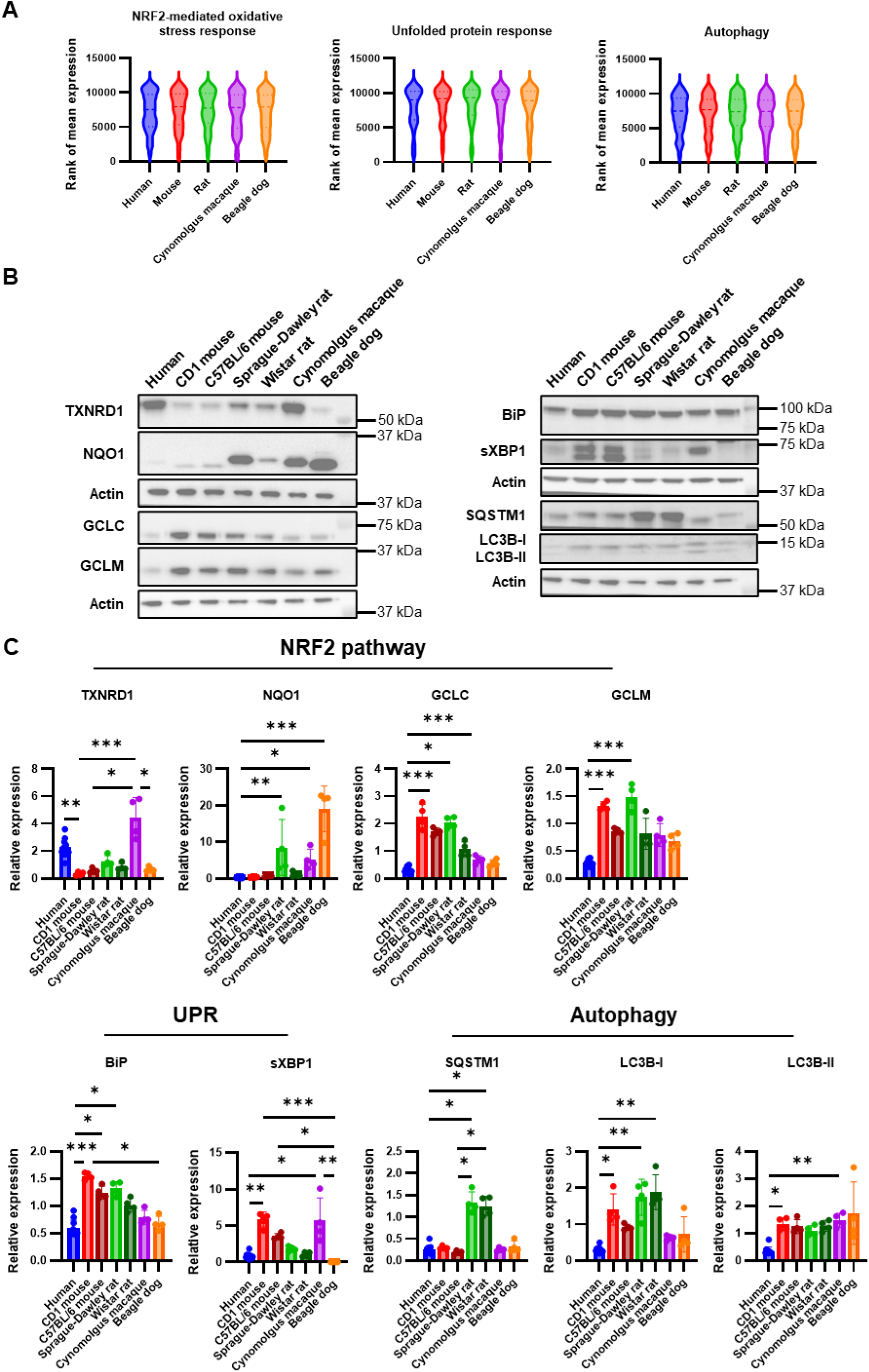
Species differences in basal expression of stress response pathway components. (A) Inter-species comparison of the rank of mean expression of genes associated with IPA canonical stress response pathways ‘NRF2-mediated oxidative stress response’ (144 genes), ‘unfolded protein response’ (69 genes) and ‘autophagy’ (167 genes) performed using a Friedman test. (B) Western blot visualisation and (C) densitometric quantification of the expression of stress response-associated proteins in the liver of humans (n=11) and various preclinical animal species (n=4/species). Target protein expression was normalised to β-actin. Data represent mean ± standard deviation. Inter-species comparisons performed using a Kruskal-Wallis test followed by Dunn’s post-hoc test. Significant inter-species comparisons indicated on the graph: * *p*_adj_ ≤ 0.05, ** *p*_adj_ ≤ 0.01, *** *p*_adj_ ≤ 0.001.

### RAT HEPATOCYTES EXHIBIT A HIGH CAPACITY FOR AUTOPHAGY

Autophagy is primarily regulated by the post-translational modification of its machinery, rather than by large-scale transcriptional adaptation as in the case of the NRF2 pathway and UPR. Therefore, western blotting was used to assess the degradation of sequestosome 1 (SQSTM1) and lipidation of microtubule-associated proteins 1A/1B light chain 3B (LC3B) in hepatocytes in the presence of the mTORC1 inhibitor, Torin1, with or without BA1, an inhibitor of lysosomal acidification and autophagosome-lysosome fusion (Figure 2A).

**Figure 2.**
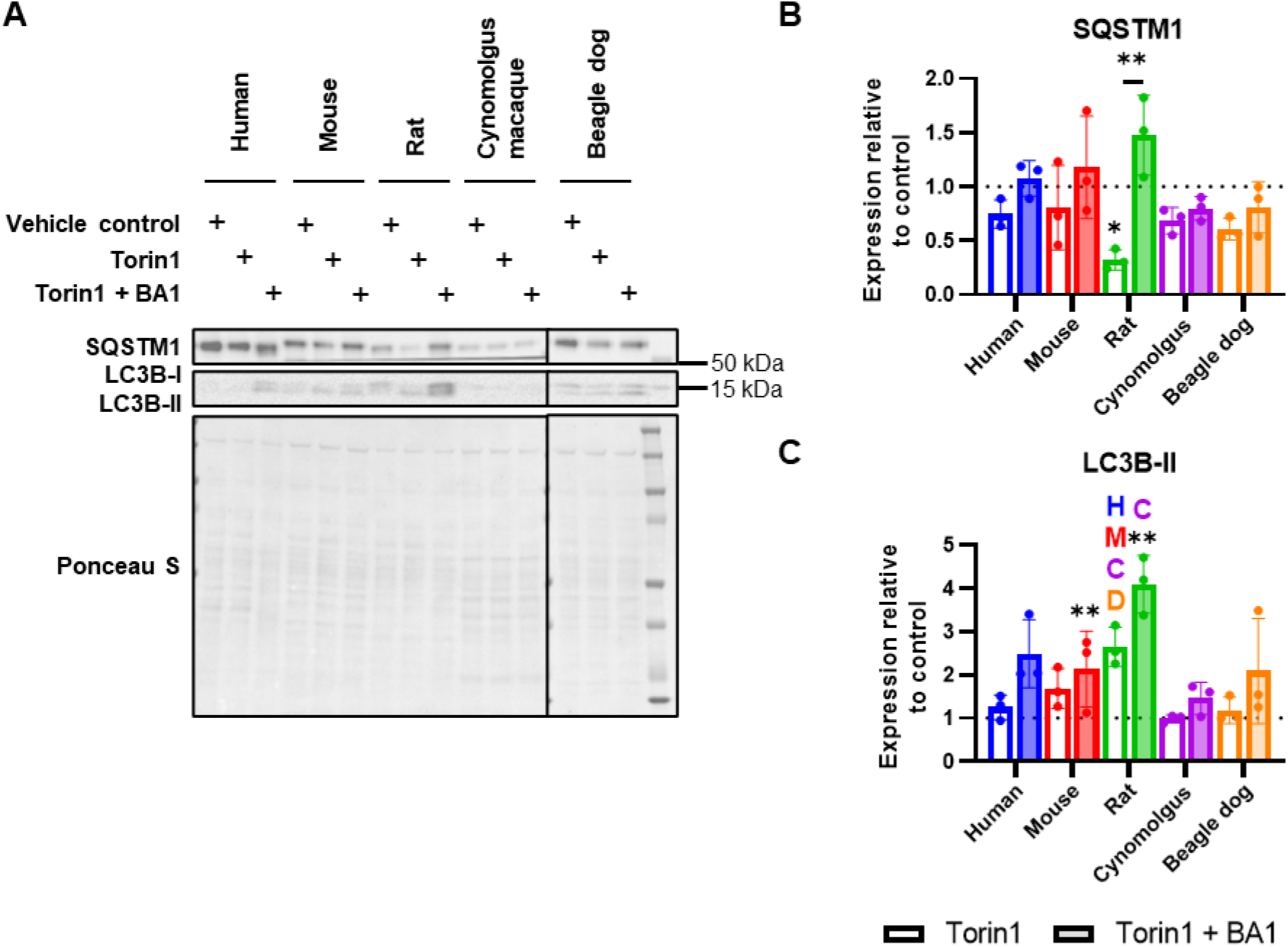
Activation of autophagy in hepatocytes from humans and important preclinical species. (A) Western blot visualisation and densitometric quantification of (B) SQSTM1 and (C) LC3B-II protein expression in primary hepatocytes following 8 h exposure to 3 µM Torin1 with or without 2 h pre-treatment and co-treatment with 30 nM Bafilomycin A1 (BA1). Target protein expression was normalised to total protein expression. Data represent mean ± standard deviation of n=3 biological replicates. Following normality assessment using a Shapiro-Wilk test, differences between treatment groups and species were assessed using a One-Way ANOVA and Tukey’s post-hoc test. Asterisks denote significant changes relative to vehicle control (floating asterisks) and between treatment conditions (asterisks on lines); * *p*_adj_ ≤ 0.05, ** *p*_adj_ ≤ 0.01. Letters denote significantly greater induction than the indicated species (*p*_adj_ ≤ 0.05): H, human; M, mouse; C, cynomolgus macaque; D, beagle dog.

While all other species displayed muted induction of autophagy, Torin1 treatment resulted in a significant decrease in SQSTM1 protein in rat hepatocytes that was wholly abolished by co-treatment of cells with the autophagic flux inhibitor, BA1, indicating an increased capacity for lysosomal degradation of autophagic cargo in this species (Figure 2B). Additionally, as shown in Figure 2C, Torin1 increased the formation of LC3B-II in rat hepatocytes to a significantly greater extent than all other species of interest. This accumulation was further enhanced by BA1 co-treatment, confirming that Torin1-induced LC3B-II elevations reflected increased autophagic flux rather than impaired degradation (Figure 2C). In all, Sprague-Dawley rat hepatocytes demonstrated the most pronounced autophagic flux *in vitro* based on both SQSTM1 degradation and LC3B-II accumulation while humans and all other preclinical species investigated exhibited relatively weak responses, which may reflect intrinsic species differences in autophagy capacity.

### RODENT HEPATOCYTES EXHIBIT A STRONG UNFOLDED PROTEIN RESPONSE

The UPR restores homeostasis following ER stress via a biphasic adaption involving an initial arrest of global translation that lessens cellular folding load followed by a transcriptional response that promotes protein folding and degradation (Reid *et al*., 2014). Therefore, to explore inter-species differences in UPR capacity, RNA-seq was employed to quantify changes in hepatocyte mRNA expression caused by the ER stress inducer, thapsigargin.

Global transcriptional adaptation to ER was stress was far weaker in primary hepatocytes from the non-rodent preclinical species investigated than in those of humans and preclinical rodent species (Figure 3A, full lists of differentially expressed genes (DEGs) provided in Supplementary File 1).

**Figure 3.**
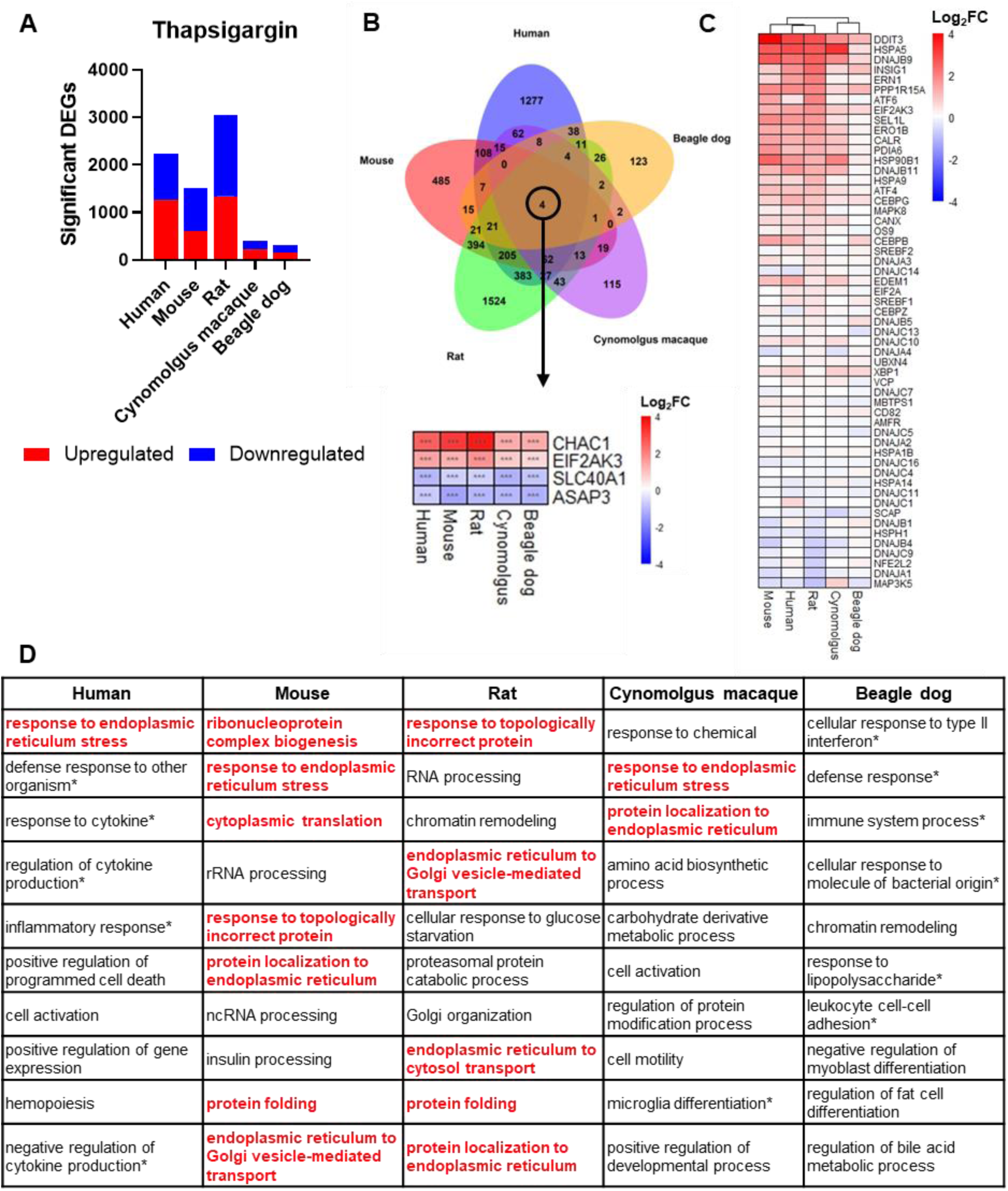
Global transcriptional effects of ER stress induction by thapsigargin. (A) Summary of the results of differential gene expression analysis for primary hepatocytes exposed to 1 µM thapsigargin for 8 h, relative to 0.5% (v/v) DMSO vehicle control (n=3/group). (B) Identification of genes differentially expressed in primary hepatocytes of all species of interest following thapsigargin exposure. Asterisks denote the significance of the change; *** *p*_adj_ ≤ 0.001. (C) Changes in expression of genes associated with the IPA canonical pathway ‘unfolded protein response’ following thapsigargin exposure, relative to vehicle control. (D) The top 10 most significantly enriched (NES ≥ 1.5, *p* ≤ 0.05) GO parent terms in each species of interest following thapsigargin exposure. Parent terms highly relevant to the UPR are highlighted. Parent terms associated with immune and inflammatory processes are denoted by asterisks.

Almost half of all DEGs were downregulated in each species, which may indicate the induction of regulated inositol-requiring enzyme 1 (IRE1)-dependent decay (RIDD), which reduces the translational and folding load of the ER by selectively degrading mRNA (Le Thomas *et al*., 2021). In all species, many DEGs were species-specific or common to just two species, and just four genes –*CHAC1*, *EIF2AK3*, *SLC40A1* and *ASAP3* – were commonly changed in all species of interest (Figure 3B).

As shown in Figure 3C, assessment of the expression of genes associated with the IPA canonical pathway ‘unfolded protein response’ also confirmed that thapsigargin induced a greater magnitude of transcriptional change in human, mouse and rat than in macaque and dog, with highly and poorly responsive species clustering separately.

GSEA revealed a large number of significantly changed biological processes in all species of interest (Supplementary File 2), which were summarised into non-redundant parent terms using rrvgo. Figure 3D summarises the 10 most enriched GO parent terms in each species of interest. No UPR-associated terms were enriched in beagle dog hepatocytes, in line with the minor transcriptional induction previously identified (Figure 3A). However, although cynomolgus macaque hepatocytes also displayed a weak global transcriptional response to ER stress induction, both ‘response to endoplasmic reticulum stress’ and ‘protein localization to endoplasmic reticulum’ were highly enriched terms in this species. In contrast, despite human hepatocytes displaying a much stronger global transcriptional adaptation than the non-rodent preclinical species, only one relevant term was within the top 10 enriched terms in this species, highlighting a divergence in the inter-species trends identified across different measures of responsiveness. Instead, in human and dog hepatocytes, many of the most enriched pathways were associated with immune and inflammatory processes, which are known to be associated with ER stress responses even under sterile conditions (Garg *et al*., 2012). However, as expected, processes associated with the ER stress response and protein synthesis, folding, and transport represented most of the top enriched biological processes in mouse and rat hepatocytes (Figure 3D). To confirm the findings of RNA-seq, targeted RT-qPCR analysis was performed to assess the induction of UPR-associated genes selected from literature (*HSPA5*, *EDEM1* and spliced *XBP1*, Supplementary Figure 2A). Different inter-species trends in transcriptional induction were identified across different UPR targets, highlighting the need for global, rather than targeted, comparisons. However, this approach also showed a greater mRNA induction in rodent hepatocytes, and a lack of transcriptional adaptation in beagle dog hepatocytes.

### RODENT HEPATOCYTES EXHIBIT A STRONG NRF2-DRIVEN OXIDATIVE STRESS RESPONSE

The cellular response to oxidative stress is primarily regulated by NRF2, a transcription factor that regulates the expression of hundreds of antioxidant and cytoprotective genes (Copple *et al*., 2019). Under basal conditions, NRF2 is sequestered in the cytosol and repressed by a homodimer of Kelch-like ECH-associated protein 1 (KEAP1) (Iso *et al*., 2016). Therefore, RNA-seq was utilised to assess transcriptional adaptation to NRF2 activation by the electrophilic inhibitor of KEAP1, CDDO-Me.

Following NRF2 activation, the rodent preclinical species investigated mounted far stronger transcriptional responses than humans and the non-rodent preclinical species of interest, as presented in Figure 4A. In all species except cynomolgus macaque (due to the very minor transcriptional response observed in this species), the majority of DEGs were species specific. Only three genes – *NQO1*, *TXNRD1* and *SRXN1* – were commonly changed in all five species of interest, although a greater change in the expression of these genes was identified in rodents than in non-rodents (Figure 4B). RT-qPCR assessment of the induction of *NQO1*, *TXNRD1* and *SRXN1* in primary hepatocytes confirmed this trend (Supplementary Figure 2B). The complete results of differential gene expression analysis are provided in Supplementary File 3.

**Figure 4.**
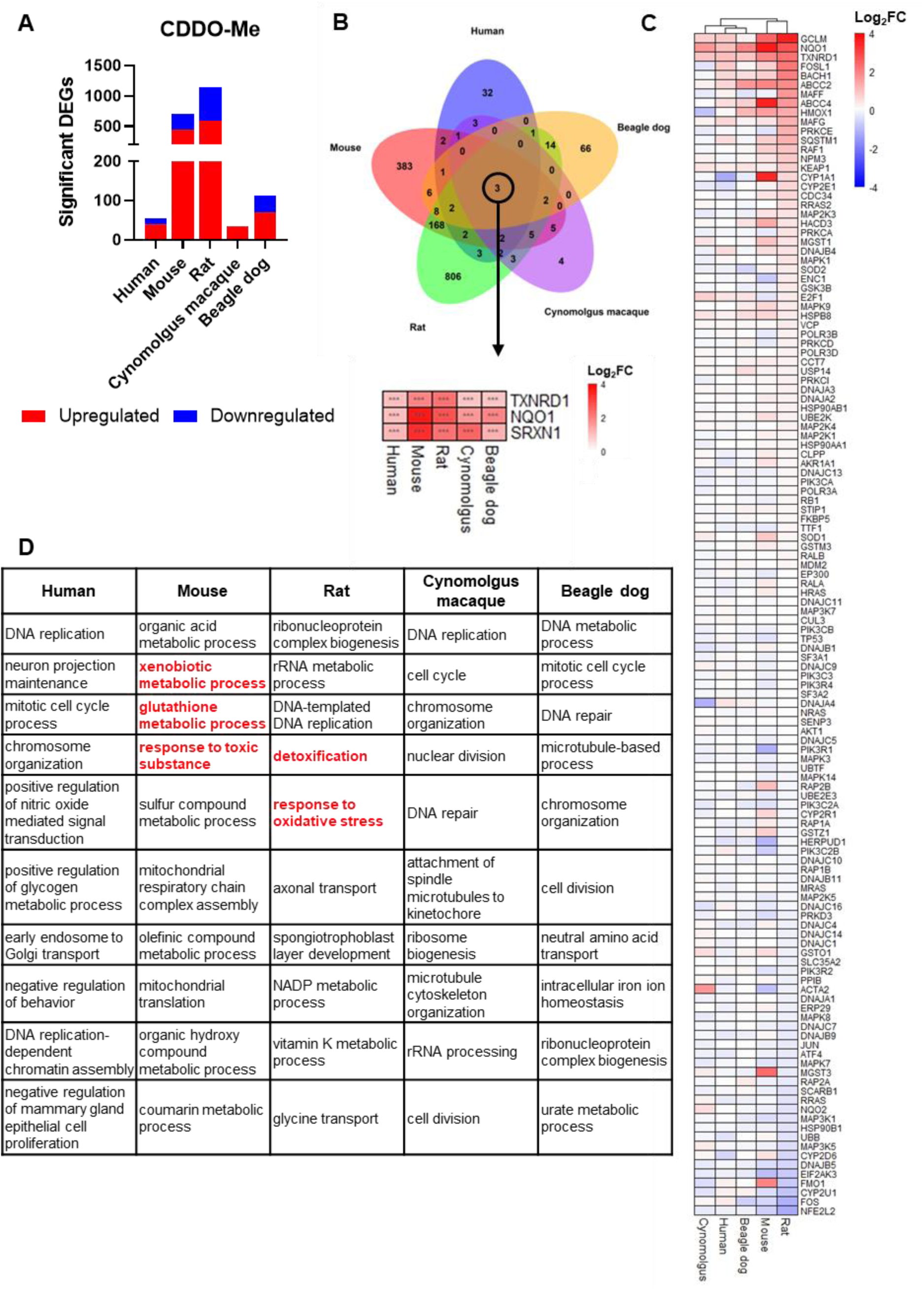
Global transcriptional effects of KEAP1 inhibition by CDDO-Me. (A) Summary of the results of differential gene expression analysis for primary hepatocytes exposed to 100 nM CDDO-Me for 24 h, relative to 0.5% (v/v) DMSO vehicle control (n=3/group). (B) Identification of genes differentially expressed in primary hepatocytes of all species of interest following CDDO-Me exposure. Asterisks denote the significance of the change; *** *p*_adj_ ≤ 0.001. (C) Changes in expression of genes associated with the IPA canonical pathway ‘NRF2-mediated oxidative stress response’ following CDDO-Me exposure, relative to vehicle control. (D) The top 10 most significantly enriched (NES ≥ 1.5, *p* ≤ 0.05) GO parent terms in each species of interest following CDDO-Me exposure. Parent terms highly relevant to the NRF2 pathway are highlighted.

Rodents also displayed a greater magnitude of change in the expression of a set of NRF2 pathway- associated genes than non-rodents, with these highly and poorly responsive species groups once again clustering separately due to their differing gene expression patterns (Figure 4C). Furthermore, as summarised in Figure 4D, GSEA identified that GO processes associated with the NRF2-mediated antioxidant response, including xenobiotic and glutathione metabolic processes, detoxification, and responses to toxic substances and oxidative stress, were among the most enriched pathways in both rodent species but were absent in humans and the non-rodent preclinical species (Supplementary File 4). Instead, processes associated with DNA replication, chromosome organisation and the cell cycle were most enriched in these species in response to CDDO-Me exposure.

Electrophilic NRF2 activators such as CDDO-Me have been shown to interact with other cellular targets, impacting pharmacological specificity (Yore *et al*., 2011). To overcome this limitation, inhibitors of the NRF2-KEAP1 protein-protein interaction (PPI) have been developed as more specific activators of the NRF2 response (Tran *et al*., 2019). Hence, to further explore species differences in NRF2 pathway activity and to evaluate whether they are conserved across different mechanisms of NRF2 activation, transcriptional responses to the PPI inhibitor, Ki696 (Davies *et al*., 2016), were investigated.

Similarly to the effects of CDDO-Me, mouse and rat hepatocytes displayed the greatest global transcriptional responses to Ki696, with weaker responses observed in human, cynomolgus macaque and beagle dog hepatocytes (Figure 5A, Supplementary File 5). Most DEGs remained species-specific, but a larger panel of nine genes was identified to be commonly changed by Ki696 across all species (Figure 5B). This panel included *NQO1*, *TXNRD1* and *SRXN1*, which were also upregulated by CDDO-Me in all species, suggesting that this subset of genes may form the most robust and translatable markers of NRF2 pathway activation, supporting previous conclusions by Morgenstern *et al*. (2024).

**Figure 5.**
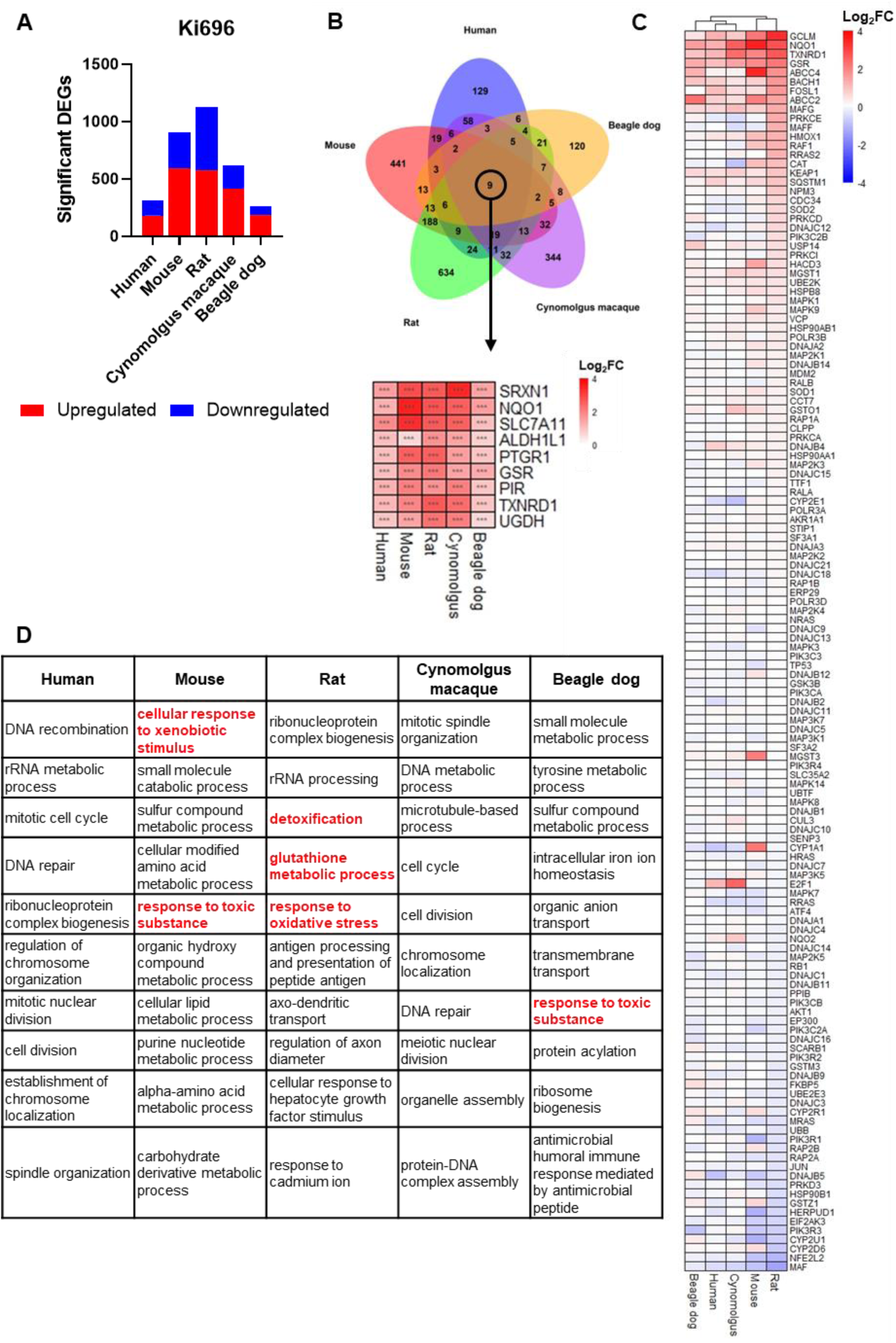
Global transcriptional effects of KEAP1 inhibition by Ki696. (A) Summary of the results of differential gene expression analysis for primary hepatocytes exposed to 10 µM Ki696 for 24 h, relative to 0.5% (v/v) DMSO vehicle control (n=3/group). (B) Identification of genes differentially expressed in primary hepatocytes of all species of interest following Ki696 exposure. Asterisks denote the significance of the change; *** *p*_adj_ ≤ 0.001. (C) Changes in expression of genes associated with the IPA canonical pathway ‘NRF2-mediated oxidative stress response’ following Ki696 exposure, relative to vehicle control. (D) The top 10 most significantly enriched (NES ≥ 1.5, *p* ≤ 0.05) GO parent terms in each species of interest following Ki696 exposure. Parent terms highly relevant to the NRF2 pathway are highlighted.

Targeted assessment of the expression of these genes also confirmed that rodent hepatocytes exhibit stronger NRF2-mediated transcriptional responses than non-rodent hepatocytes (Supplementary Figure 2C). Most of the remaining commonly changed DEGs (*SLC7A11*, *PTGR1*, *GSR*, *PIR* and *UGDH*) were differentially expressed in multiple, but not all, species of interest following CDDO-Me treatment. Also in line with responses to CDDO-Me, rodents displayed a greater activation of the NRF2 pathway by Ki696 compared to non-rodents, as measured by changes in the expression of NRF2-associated genes (Figure 5C, Supplementary File 6) and the enrichment of NRF2-associated biological processes in GSEA (Figure 5D). Furthermore, Ki696 caused the enrichment of GO processes associated with DNA replication and the cell cycle in human and cynomolgus macaque hepatocytes, indicating that the activation of these pathways is associated with NRF2 signalling in these *in vitro* models, as opposed to the off-target effects of CDDO-Me.

## Discussion

The detection of hepatotoxicity liabilities during preclinical studies is essential in minimising patient risk during clinical trials and post-approval use. Until recently, most major drug regulators mandated the use of *in vivo* animal studies in the preclinical safety investigation of a new small molecule drug. However, animal studies poorly predict human DILI (Olson *et al*., 2000, Tamaki *et al*., 2013) due to an incomplete understanding of the intrinsic similarities and differences between humans and preclinical model species and the impact on a species’ relative sensitivity to toxicity. As sufficiently predictive and validated animal-free new approach methodologies (NAMs) are not yet approved, animal studies will likely remain a part of drug safety evaluation for the foreseeable future (Madorran *et al*., 2020). Hence, there remains a need to improve our understanding of the differences between humans and preclinical species to improve the translatability and predictivity of animal toxicology studies. This study compared the capacities of key DILI-associated stress response pathways in humans and preclinical species. Our findings highlight notable inter-species differences that should be considered during preclinical species selection and the interpretation of preclinical toxicology studies.

We recently reported the effect of differential stress response pathway activation in laboratory mice and rats on the hepatotoxicity of APAP (Russomanno *et al*., 2023), which exerts its toxicity via a CRM (Corcoran *et al*., 1980, Zhang *et al*., 2022). Our findings were consistent with the work of Xu *et al*. (2021), who investigated the differential responses of mice and rats to the hepatotoxin psoraleae fructus (FR). Importantly, we accounted for species-specific differences in APAP bioactivation via the determination of APAP doses that generated equivalent CRM burden in each species (Russomanno *et al*., 2023). In the present study, we expanded the number of species considered and utilised specific, positive-control pharmacological modulators of the relevant pathways to assess stress response activation *in vitro* without confounding factors associated with species-specific bioactivation of a hepatotoxin or non-specific interactions of a CRM.

We have found that the rodent preclinical species investigated – particularly rats – display a notably greater hepatic capacity for adaptation to stress response activation than humans and the most commonly used non-rodent preclinical species, cynomolgus macaques and beagle dogs. In a review of the preclinical and clinical safety of approved drugs, Olson *et al*. (2000) found that non-rodent preclinical species predict human toxicities far more accurately than rodent preclinical species. The similar basal and adaptive stress response capacities of non-rodent preclinical species and humans identified in this study may contribute to these greater predictive capabilities. In contrast, the robust stress response capacity of rodents – particularly rats – identified in this study may render these species less sensitive than humans to hepatic chemical insult and DILI, contributing to the relatively poor detection of human toxicities by these species (Olson *et al*., 2000, Monticello *et al*., 2017). In the future, the scope of our work can be expanded by considering other stress responses implicated in DILI, such as the DNA damage and mitochondrial respiratory responses, as well as other tissues relevant to common drug toxicities. In addition, further mechanistic work is necessary to correlate stress response capacity to DILI sensitivity in relevant species.

Many hepatotoxins exert their toxicity through CRMs, which activate cellular stress responses, making stress-response markers attractive as early indicators of CRM-mediated DILI in preclinical studies (Monroe et al., 2020; Guo and van den Beucken, 2024). However, the translatability of transcriptional biomarker panels across preclinical models and species remains unclear. Morgenstern et al. (2024) identified six NRF2 activity markers conserved in humans, mice, and rats – *NQO1*, *TXNRD1*, *SRXN1*, *HMOX1*, *GCLC*, *GCLM*. Our findings further support the robustness of three of these genes (*NQO1*, *TXNRD1*, *SRXN1*) as translatable NRF2 biomarkers across a wider range of preclinical species. In contrast, no similar attempts have been made to establish a translatable panel of UPR activity biomarkers. Fei et al. (2025) reported the use of a panel of UPR-related genes as prognostic biomarkers in acute myeloid leukaemia. However, aberrant activation of the UPR in leukaemia may render this panel inappropriate for the prediction of DILI potential in normal cells, whilst the wider species relevance of the panel is unclear. To address this gap, we leveraged RNA-seq data to identify four genes (*EIF2AK2*, *CHAC1*, *SLC40A1*, and *ASAP3*) commonly changed across humans and all preclinical species, representing a candidate panel of translatable UPR markers. Although further validation is required to assess the suitability of these biomarkers in relevant preclinical models, such panels may prove valuable in early hepatocyte-based *in vitro* screens for CRM liability in new drug candidates.

In all, this study highlights notable intrinsic species-specific differences in a factor known to influence the progression of DILI, which should be accounted for during species selection to maximise the translatability of preclinical toxicology studies. Consideration of the enhanced hepatic capacity of rodents to mitigate cellular stress and chemical insult, which may underlie a relative resistance of these species to CRM-mediated DILI (Xu *et al*., 2021, Russomanno *et al*., 2023), could contribute to improving the predictive capabilities of preclinical *in vivo* toxicology studies. However, as a complete move away from poorly predictive rodents to more predictive non-rodent species is infeasible due to ethical and practical considerations that must be made during species selection (Son *et al*., 2020), our findings also reinforce the existing need to reduce reliance on animal models – which will always possess the potential to display species-specific differences to humans – in drug safety evaluation.

This transition is encouraged by amendments to legislation and regulatory guidance, such as the US Food and Drug Administration (FDA) Modernization Act 2.0 (2022), which eliminated the legal requirement for animal toxicology studies to support new drug applications submitted to the FDA.

The aspiration is that, as animal-free NAMs are increasingly validated and integrated into regulatory frameworks, they will ultimately replace traditional *in vivo* animal toxicology studies, providing more human-relevant, predictive approaches for evaluating drug safety during preclinical development.

---

**Supplementary Figure 1.**
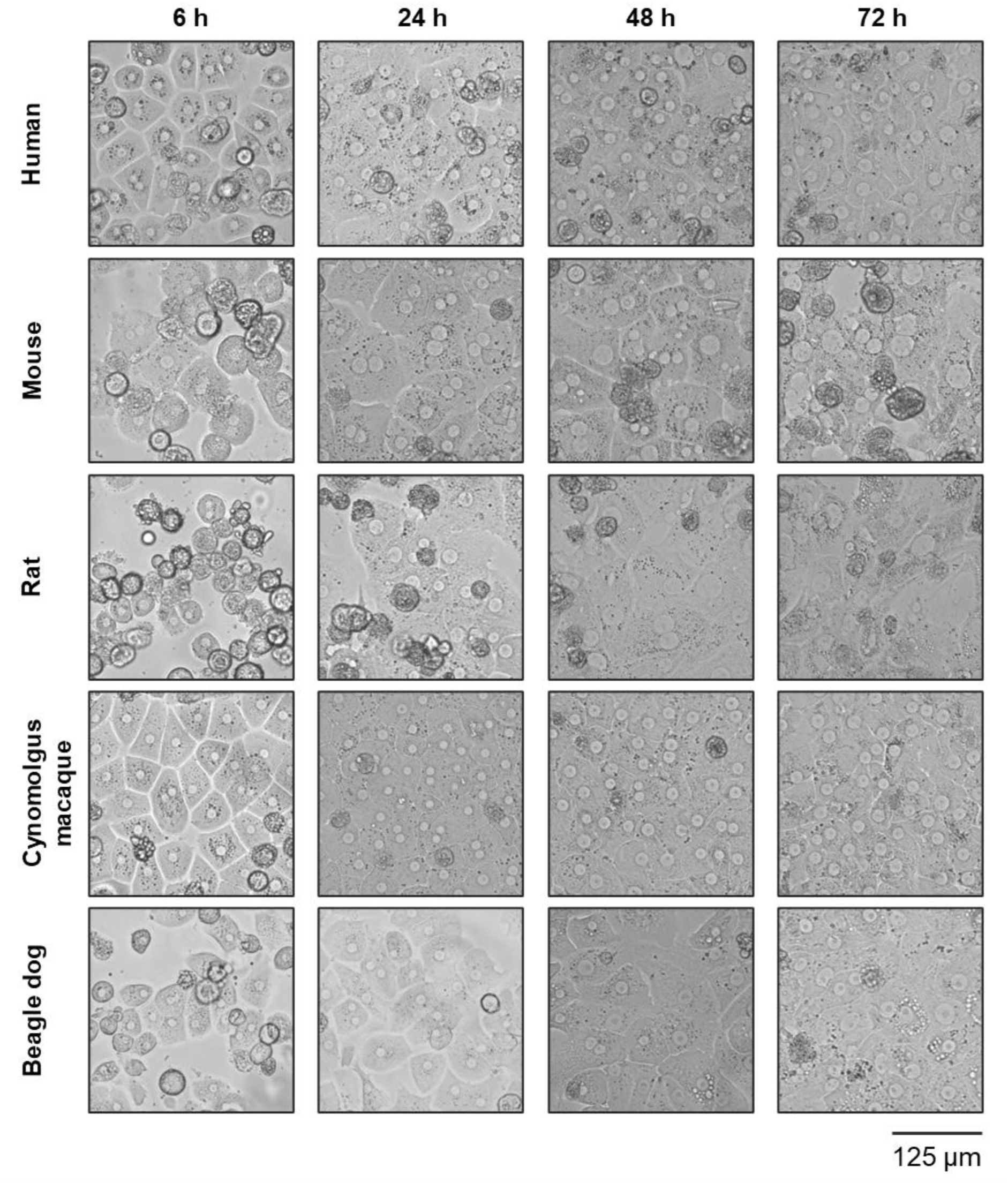
Primary hepatocytes cultured in serum-supplemented medium retain their morphology for at least 48 h in vitro. Morphology of human, CD1 mouse, Sprague-Dawley rat, cynomolgus macaque and beagle dog primary hepatocytes cultured in serum-supplemented hepatocyte culture medium 6, 24, 48 and 72 h post-plating in 2D monolayer culture on a collagen I-coated plated. Scale bar shown represents 125 µm.

**Supplementary Figure 2.**
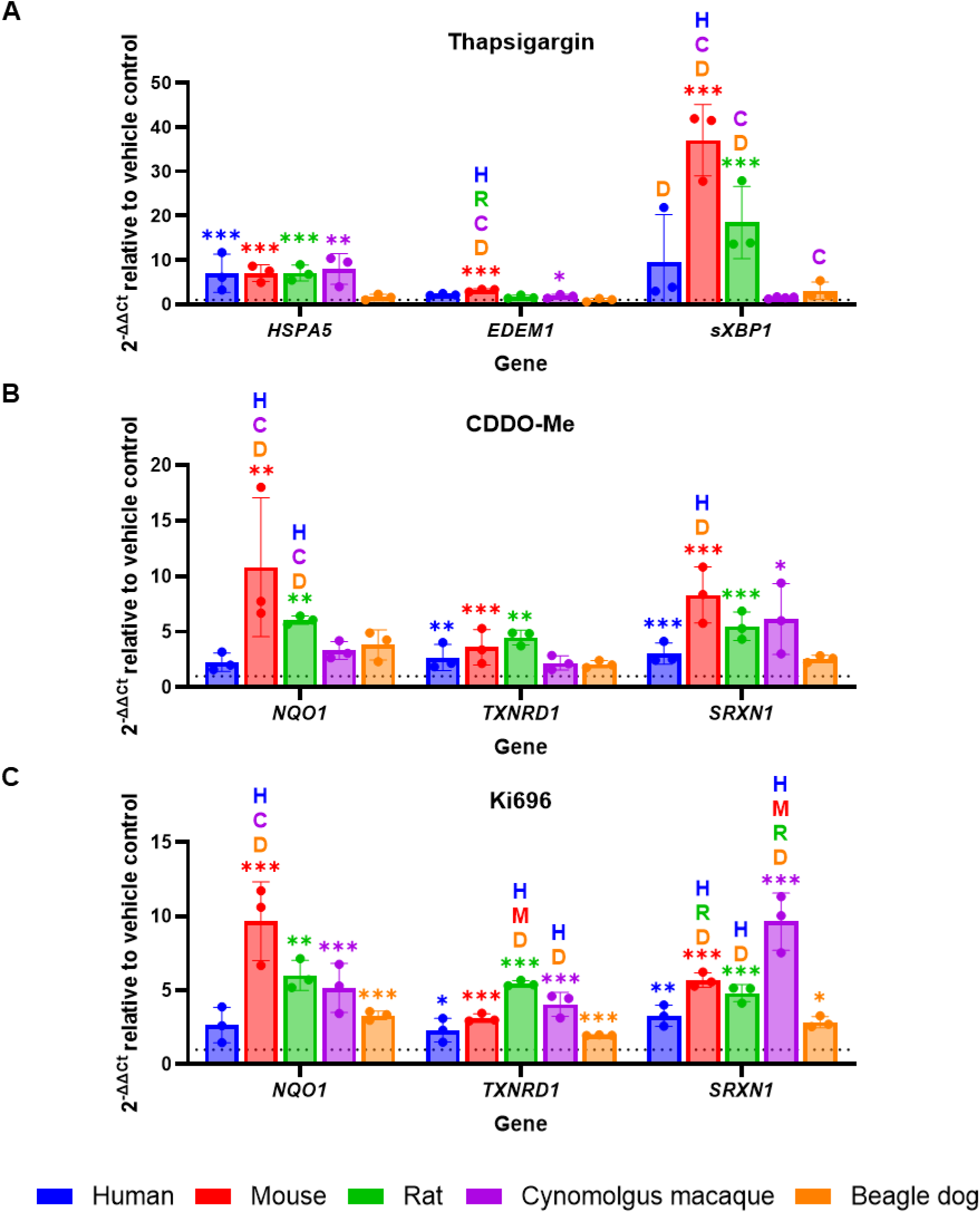
Targeted assessment of stress response induction. Induction of mRNA targets of (A) the UPR or (B & C) the NRF2 pathway in primary hepatocytes by (A) thapsigargin (1 μM, 8 h), (B) CDDO-Me (100 nM, 24 h), or (C) Ki696 (10 μM, 24 h), measured by RT-qPCR. All data represent mean ± standard deviation of three biological replicates (n=3) of 2^-ΔΔCt^ of target gene relative to vehicle control and normalised against expression of the housekeeping genes GAPDH and ACTB. Significant differences relative to vehicle control and between species assessed using a One-Way ANOVA and Tukey’s post-hoc test or Kruskal-Wallis and Dunn’s post-hoc test, as appropriate. Asterisks denote significant changes relative to vehicle control; * p_adj_ ≤ 0.05, ** p_adj_ ≤ 0.01; *** p_adj_ ≤ 0.001. Letters denote significantly greater induction than the indicated species (p_adj_ ≤ 0.05): H, human; M, mouse; R, rat; C, cynomolgus macaque; D, beagle dog.

**Supplementary Table 1.**
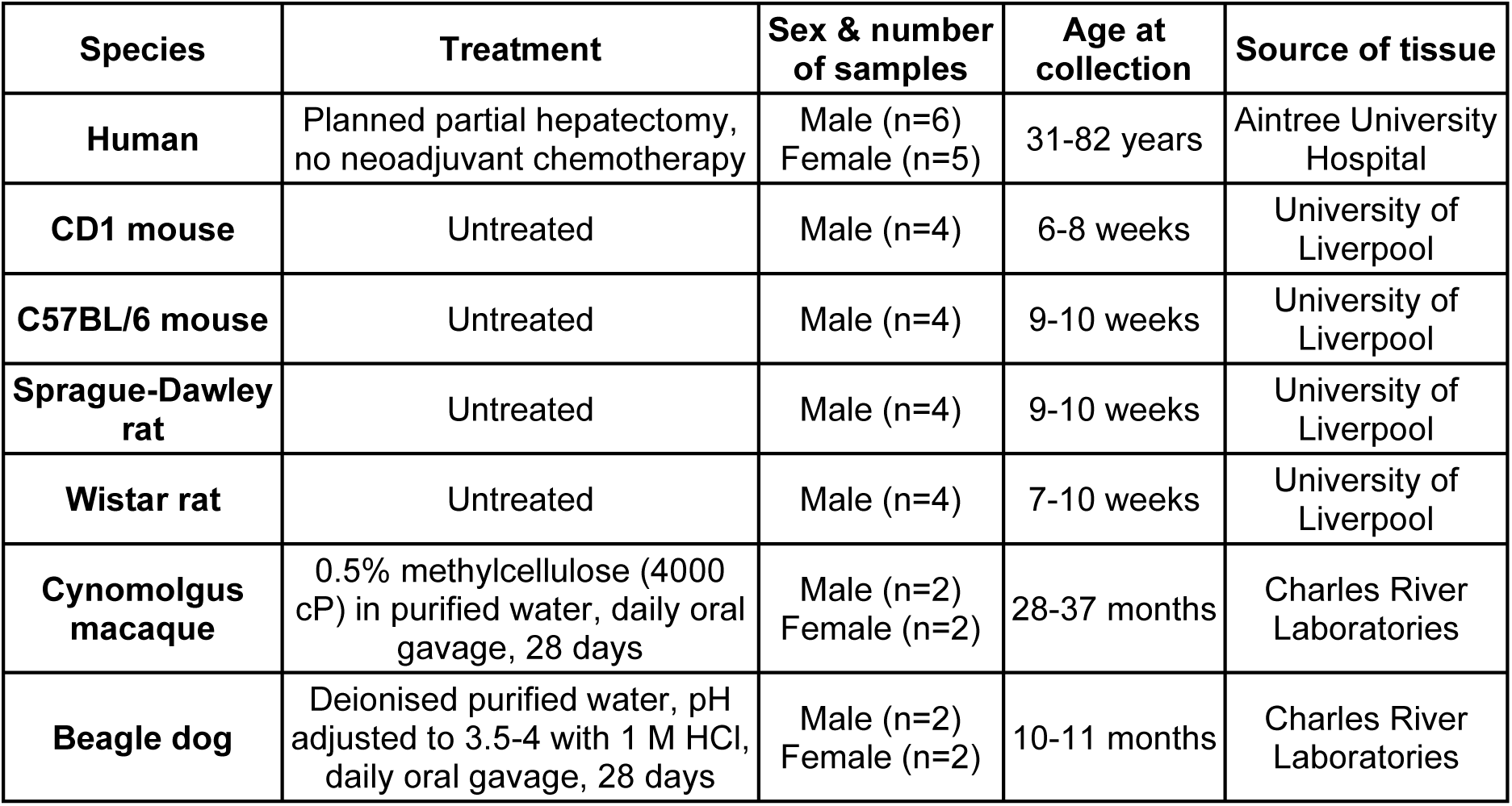
Liver tissue sample information.

**Supplementary Table 2.** Western blot antibody information.

| Protein | Antibody supplier & code | Molecular weight (kDa) | Dilution | Host species |
| --- | --- | --- | --- | --- |
| β-Actin | Abcam (ab6276) | 42 | 1:10,000 | Mouse |
| TXNRD1 | Abcam (ab124954) | 55 | 1:5000 | Rabbit |
| NQO1 | Abcam (ab2346) | 30 | 1:2000 | Goat |
| GCLC | Abcam (ab41463) | 73 | 1:5000 | Rabbit |
| GCLM | Abcam (ab126704) | 31 | 1:5000 | Rabbit |
| BiP | Proteintech (11587-1-AP) | 78 | 1:5000 | Rabbit |
| sXBP1 | Proteintech (24868-1-AP) | 70 | 1:2000 | Rabbit |
| SQSTM1 | Sigma (P0067) | 62 | 1:2500 | Rabbit |
| LC3B | Cell Signaling Technology (2775) | 14 & 16 | 1:750 | Rabbit |
| HRP-conjugated anti-mouse IgG | Sigma-Aldrich (A9044) | NA | 1:5,000 | NA |
| HRP-conjugated anti-rabbit IgG | Sigma-Aldrich (A9169) | NA | 1:5,000 | NA |
| HRP-conjugated anti-goat IgG | Dako (P0449) | NA | 1:5,000 | NA |

HRP, horseradish peroxidase.

**Supplementary Table 3.**
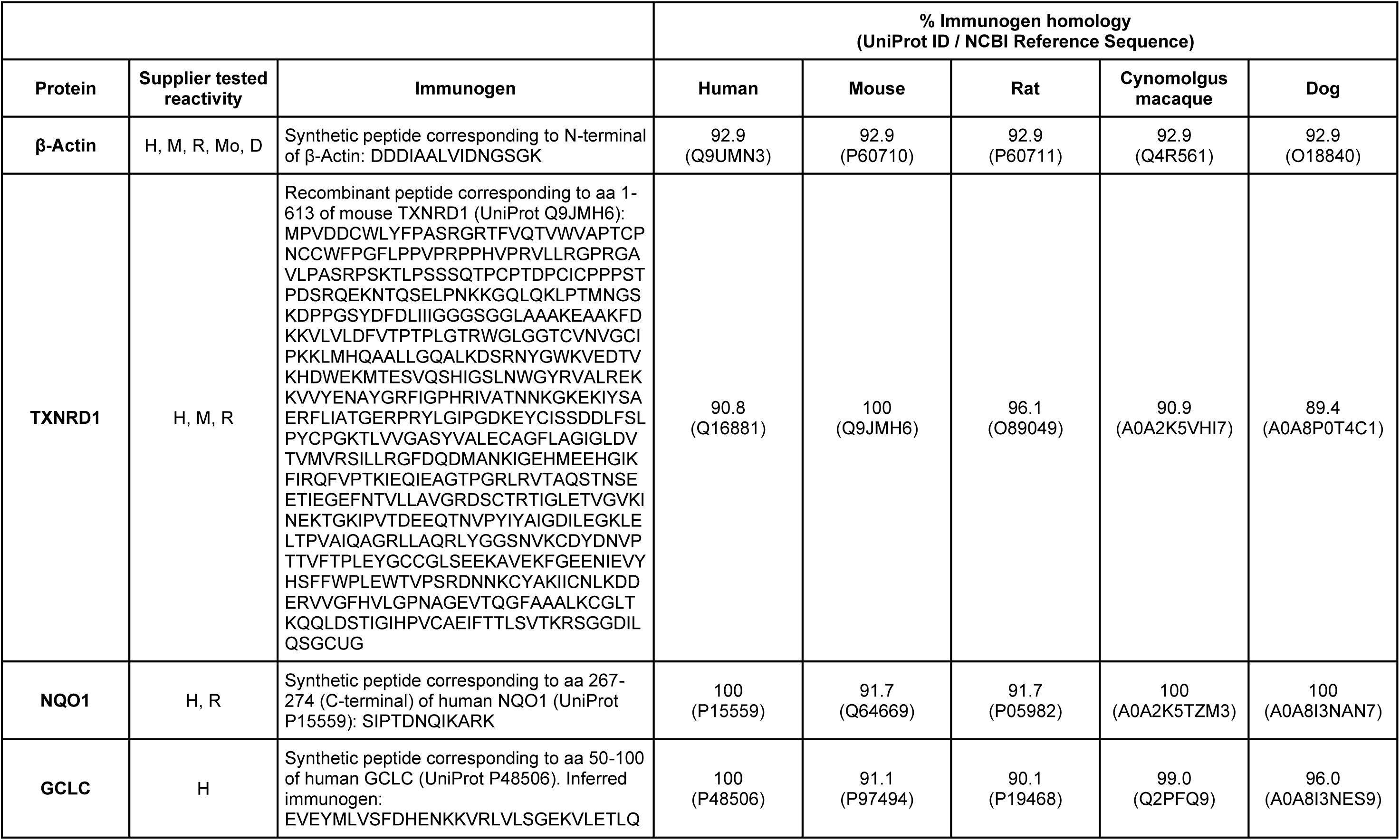
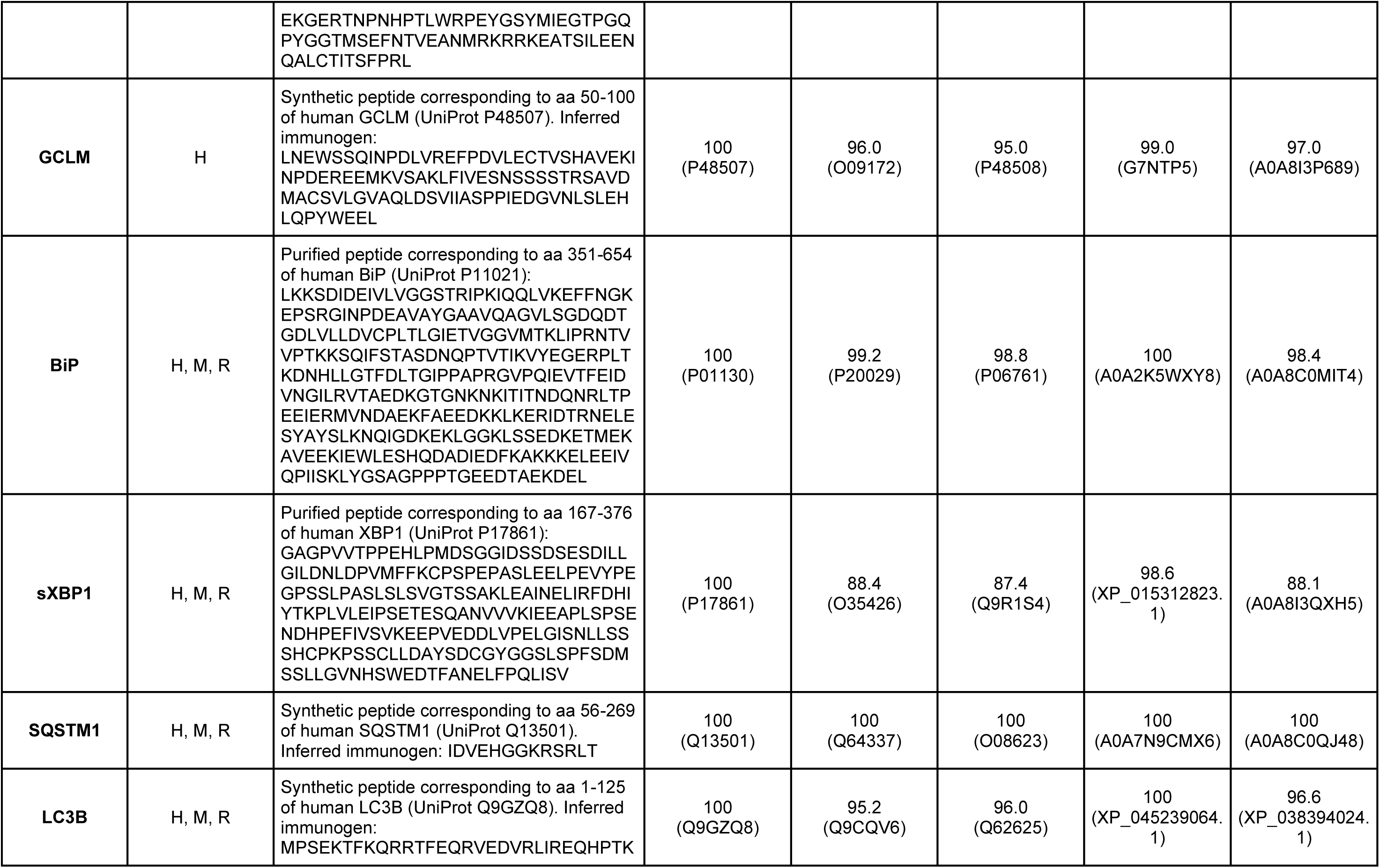
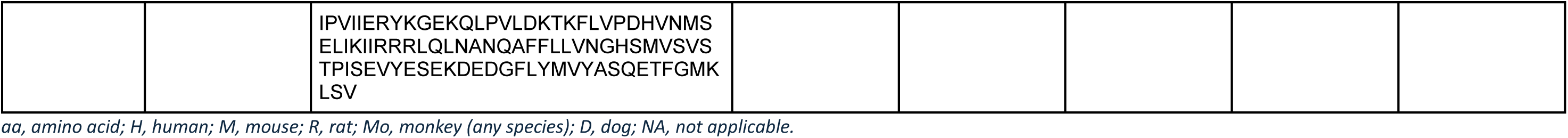
Primary antibody immunogen homology across species of interest.

